# Ultraviolet screening by slug tissue and tight packing of plastids protect photosynthetic sea slugs from photoinhibition

**DOI:** 10.1101/2021.07.15.452583

**Authors:** Vesa Havurinne, Riina Aitokari, Heta Mattila, Ville Käpylä, Esa Tyystjärvi

## Abstract

- One of the main unsolved questions regarding photosynthetic sea slugs is how the slug plastids handle photoinhibition of Photosystem II. Photoinhibition has not been studied in detail in these animals although resilience against photoinhibition might obviously explain the longevity of plastids inside animal cytosol.
- Light response and action spectrum of photoinhibition were measured from the slug *Elysia timida* and its prey alga *Acetabularia acetabulum*. Plastid packing in the slugs and algae was compared with spectroscopic and microscopic methods. The importance of plastid concentration was also estimated by measuring photoinhibition from starved slugs.
- Compared to *A. acetabulum, E. timida* is highly resistant against photoinhibition. The resilience of the slugs is even more pronounced in the UV-region, as the slug tissue screens UV radiation. The plastids in the slug tissue are tightly packed, and the outer plastids protect the inner ones from photoinhibition.
- The sea slug *E. timida* protects its plastids from photoinhibition by screening UV radiation and packing the plastids tightly in its tissues. Both mechanisms enhance the longevity of the plastids in slug cytosol and ameliorate the need for repair of photoinhibited Photosystem II.

## Introduction

*Elysia timida* belongs to a group of animals that carry out photosynthesis using plastids stolen from their prey. This interesting phenomenon, called kleptoplasty, has only been reported in Sacoglossan sea slugs like *E. timida* (Rumpho *et al*., 2011; de Vries *et al*., 2014) and marine flatworms (Van Steenkiste *et al*., 2019). The record holding photosynthetic slug, *E. chlorotica*, maintains kleptoplasts functional for approximately a year (Green *et al*., 2000), and has served as one of the most important subjects for the study of kleptoplastic animals (Chan *et al*., 2018; Cai *et al*., 2019). However, the limited availability of *E. chlorotica* individuals is a major obstacle for in-depth laboratory studies. Plastids of the slug *E. timida* originate from the green alga *Acetabularia acetabulum* (hereafter *Acetabularia*). *Elysia timida* is known for its easy husbandry in the laboratory (Schmitt *et al*., 2014; Havurinne & Tyystjärvi, 2020). Use of laboratory cultures reduces stress on natural populations of sea slugs and offers controlled conditions that improve the reproducibility of the experiments.

Many questions related to photosynthetic sea slugs have no answer so far. For example, it is unclear how the slugs recognize and incorporate foreign organelles into their own cells. The uptake process has been suggested to involve the slug’s innate immune system that can possibly recognize the plastids via scavenger receptors and thrombospondin-type-1 repeat proteins (Chan *et al*., 2018; Clavijo *et al*., 2020). It is also uncertain just how important are the native properties of the plastids themselves in terms of facilitating their survival for weeks and months inside animal cytosol in isolation from the algal nucleus. The slugs are only able to retain plastids that come from specific algae species (Christa *et al*., 2013; de Vries *et al*., 2013), but to what extent this is due to the general robustness of plastids of these algae (Giles & Sarafis, 1972; Trench *et al*., 1973a; 1973b; Green *et al*., 2005) or their specific genetic and photosynthetic properties (de Vries *et al*., 2013; Christa *et al*., 2018; Havurinne *et al*., 2021) remains to be fully tested.

Irreversible light-induced damage to Photosystem II (PSII) of the photosynthetic electron transfer chain, termed photoinhibition, is an important reason why survival of plastids in isolation within slug cells for months requires special mechanisms. Photoinhibition, the paradoxical downside of utilizing light energy to run photosynthesis, has been shown to be ubiquitous in photosynthetic organisms, and it occurs even in low light (Tyystjärvi & Aro, 1996). Even though the exact mechanism(s) of photoinhibition remain elusive, decades of work on the topic have revealed several “rules” that most photosynthetic organisms comply to. These rules include: (I) direct proportionality of the rate constant of the damaging reaction with photosynthetic photon flux density (PPFD), (II) the damaging reaction proceeds according to first-order reaction kinetics, and (III) UV radiation is considerably more damaging than visible light (Tyystjärvi, 2013). In spite of photoinhibition, photosynthetic organisms maintain high photosynthetic activity by continuously repairing damaged PSII reaction centers (Järvi *et al*., 2015). Because the repair cycle of PSII is efficient, the actual rate of photodamage to PSII can be measured only if the repair cycle is blocked with an antibiotic that blocks plastid translation, such as lincomycin.

Even though resilience against the damaging reaction of photoinhibition might explain the longevity of plastids inside photosynthetic sea slugs, very few studies have addressed this directly. The effect of different intensities of light on the longevity of PSII activity in the plastids of *E. timida* and *E. viridis* have been evaluated, and the results show that stronger light during starvation leads to shorter retention of functional plastids (Vieira *et al*., 2009; Christa *et al*., 2018). On the other hand, much effort has been invested into evaluating the physiological photoprotection mechanisms of the plastids. It has been shown that the plastids in the slugs maintain similar or slightly elevated photoprotective non-photochemical quenching (NPQ) mechanisms as the plastids in the algae, at least in recently fed slugs (Cruz *et al*., 2015; Christa *et al*., 2018; Cartaxana *et al*., 2019; Havurinne & Tyystjärvi, 2020). However, the effectiveness of these NPQ mechanisms in preventing net photoinhibition in *E. timida* in the absence of lincomycin remains controversial (Christa *et al*., 2018; Cartaxana *et al*., 2019). Photoinhibition and subsequent recovery of PSII in photosynthetic sea slugs in the presence of lincomycin has only been evaluated twice; once by Christa *et al*., (2018) in *E. timida* and *E. viridis*, and recently by us in *E. timida* (Havurinne *et al*., 2021). The results of these two studies contradict each other, as Christa *et al*. did not find any significant differences in the recovery of PSII in the presence or absence of lincomycin, whereas our data showed that lincomycin efficiently blocks the recovery process in photoinhibited plastids of *E. timida*. This clearly emphasizes the need for further studies into both the actual damaging reactions and recovery processes from photoinhibition in photosynthetic sea slugs.

Here, we set out to thoroughly examine the characteristics of photoinhibition in the sea slug *E. timida* and its prey green alga *Acetabularia*, with the idea of testing which of the rules of photoinhibition hold true in the slugs. While our photoinhibition experiments show that the slugs are governed by the same basic principles of photoinhibition as their algal counterparts, properties of the slug tissue and placement of the plastids inside slug cells drastically slow down photoinhibition of the plastids inside the slug *E. timida*.

## Materials and methods

### Organisms and culture conditions

The sea slug *E. timida* (strain TI1) and its prey green alga *Acetabularia* (strain DI1; originally isolated by Diedrik Menzel) were maintained in 10 l plastic tanks at 23 °C in a 12/12h day/night cycle (PPFD 40-50 µmol m^-2^ s^-1^ during the day), as described earlier (Havurinne & Tyystjärvi, 2020). *Elysia timida* was cultured in 3.7% (m/v) artificial sea water (ASW; Sea Salt Classic; Tropic Marin, Montague, MA, USA). *Acetabularia* was cultured in f/2 medium made into 3.7% ASW. The slug tanks were continuously aerated. Experiments with *E. timida* were mainly done with freshly fed individuals, but slugs that had been kept in starvation (removed from the algal food source) for different time periods in their growth conditions were used in certain experiments, as indicated in the text. For the starvation treatments, the slugs were deprived of their food, and kept in 5 l tanks filled with ASW. The starving slugs were moved to clean tanks with fresh ASW weekly.

### Photoinhibition treatments

*Elysia timida* individuals of similar size and green color were selected for the photoinhibition treatments and subjected to overnight darkness in ASW containing 10 mg/ml lincomycin, a translation inhibitor shown to be plastid specific in plants (Mulo *et al*., 2003). *Acetabularia* samples selected for photoinhibition treatments were treated in an identical manner to the slugs in f/2 medium in the absence or presence of 10 mg/ml lincomycin. Lincomycin concentrations that have been used to block the PSII repair cycle in the plastids of photosynthetic sea slugs are high, for example 8 mg/ml (Christa *et al*., 2018) or 10 mg/ml (Havurinne *et al*., 2021). We opted to use the same concentration of lincomycin for the slugs and the algae because preliminary experiments showed that even in the algae a high lincomycin concentration is required to block PSII recovery after photoinhibition (Supporting Information Fig. S1). Effectiveness of 10 mg/ml lincomycin in stopping PSII repair cycle in the slug plastids has been illustrated previously (Havurinne *et al*., 2021).

**Supporting information figure S1.**
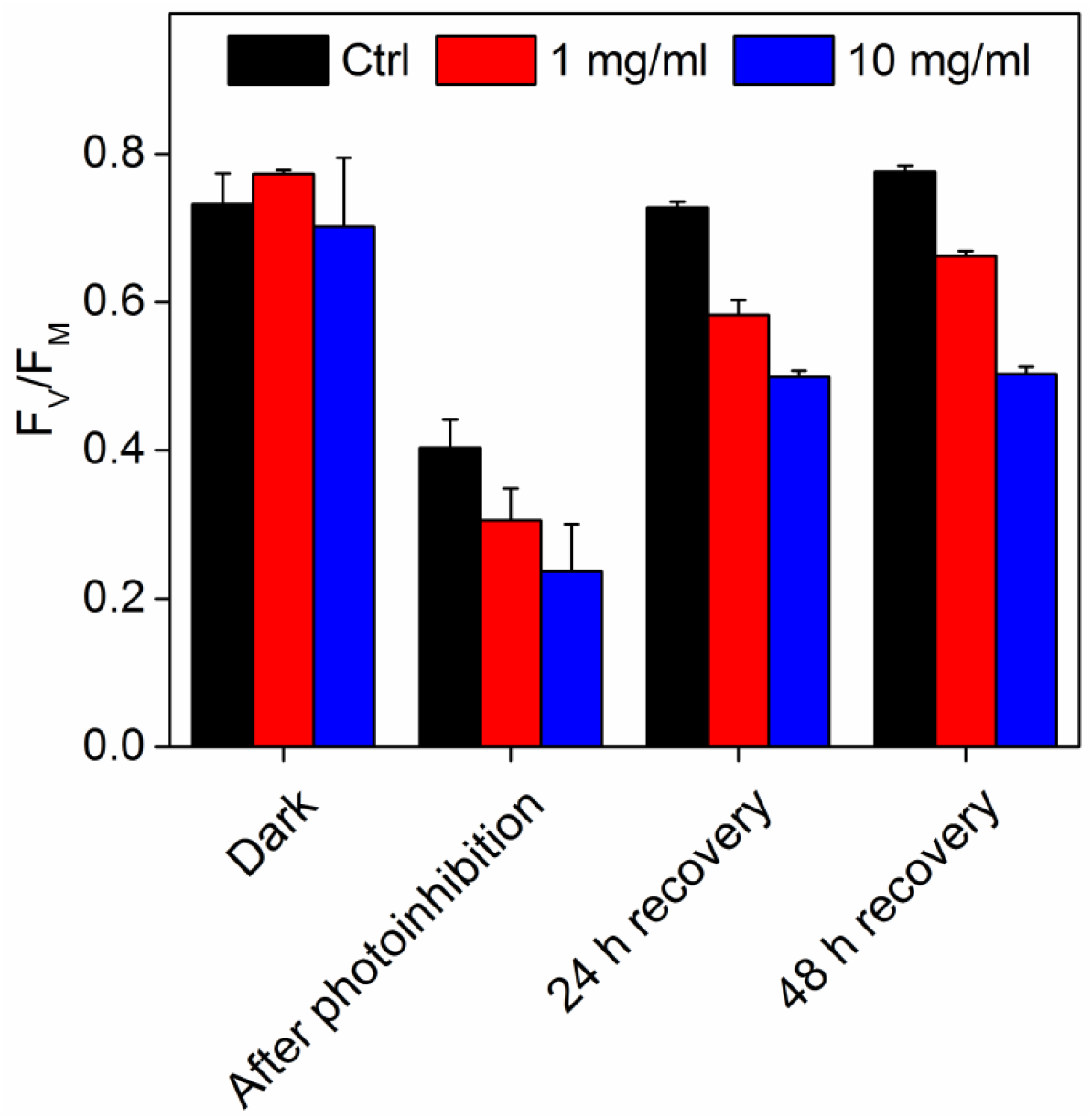
The effect of different concentrations of lincomycin on recovery of PSII activity in *Acetabularia*. All samples were kept in the dark overnight in the absence (control, black bars) or presence of 1 mg/ml (red bars) or 10 mg/ml (blue bars) of lincomycin in f/2 culture medium prior to exposing them to a 60 min high light treatment (PPFD 2000 µmol m^-2^ s^-1^). The same samples were subsequently incubated in the presence of lincomycin in the growth conditions under low light (PPFD ∼10 µmol m^-2^ s^-1^) to measure recovery from photoinhibition. The fluorescence parameter F_V_/F_M_ was used as a proxy of PSII activity, and it was measured from the samples after a minimum of 20 min dark period. Each bar represents an average of three biological replicates and the error bars show SD.

After an overnight incubation in the dark with lincomycin, slugs or algae were placed inside the wells of a 24-well plate in their respective medium. For most of the experiments one slug individual or 3-5 strands of algae were placed inside a single well of the well plate, representing one biological replicate. The well plate bottom for both species was covered with aluminum foil to ensure that the slugs receive as much light as possible, as they tend to curl up next to the edges of the wells when exposed to high light. The wells were large enough to prevent the algal strands from shading each other in the same well. The ratio of variable to maximum chlorophyll *a* fluorescence (F_V_ /F_M_) was measured with a pulse amplitude modulation fluorometer PAM-2000 (Walz, Effeltrich, Germany), and used as a proxy of PSII activity as described earlier (Havurinne & Tyystjärvi, 2020). The F_V_ /F_M_ parameter is known to function well as a probe of photoinhibitory damage (Tyystjärvi, 2013) although it was recently shown that F_V_ /F_M_ does not represent the maximum quantum yield of PSII (Sipka *et al*., 2021). The first F_V_ /F_M_ value was measured from samples that had been dark acclimated overnight, and 20 min dark incubation was applied before F_V_ /F_M_ measurements during the photoinhibition treatments. The samples were returned to light treatment thereafter. The rate constant of photoinhibition (k_PI_) was determined by fitting the decrease in F_V_ /F_M_ to first order decay kinetics (Tyystjärvi & Aro, 1996) using SigmaPlot v.14.0 (Systat Software, Inc., San Jose, CA, USA); time was measured as the cumulative illumination time, excluding the 20-min dark incubations.

White light for the photoinhibition treatments was provided by an Artificial Sunlight Module (SLHolland, Breda, The Netherlands; see Supporting Information Fig. S2 for the irradiance spectrum). The action spectra of photoinhibition were measured by exposing the samples to monochromatic light of different wavelengths. Although the emission spectra of the light sources were wide in some cases, the visible spectrum light treatments will be referred to as 690, 660, 560, 470 and 425 nm and those of the UV spectrum as 365 (UVA), 312 (UVB) and 254 nm (UVC). Monochromatic visible light used in the photoinhibition experiments was obtained using a custom-built LED array equipped with one of the Andover Corporation line filters 690FS, 660FS, 560FS and 470FS (Newport, Irvine, CA, USA), where numbers stand for the respective center wavelengths of the filters; the half width at half maximum of these filters is 10 nm. 425 nm light was obtained using the Artificial Sunlight Module (SLHolland) in combination with 450 nm short pass and 400 nm long pass filters (Newport Corporation, Franklin, MA, USA), and the UV sources were VL-8.LC (UVA and UVC) and VL-8.M (UVB) UV lamps (Vilber, Marne la Vallée, France). PPFDs (or photon flux density, PFD, for UV radiation) of the photoinhibition treatments were measured from the water surface levels of the 24-well plates either with a planar, wavelength calibrated light sensor (LI-COR Biosciences; Lincoln, NE, USA) or with an STS-UV/visible light spectrometer (Ocean Optics, Largo, FL, USA). The PPFDs of the visible light wavelengths 690, 660, 560, 470 and 425 nm were 300, 309, 134, 233 and 200 µmol m^-2^ s^-1^, whereas UVA, UVB and UVC treatments were done with the PFDs of 33, 51 and 23 µmol m^-2^ s^-1^, respectively. In order to facilitate comparison, original k_PI_ values from the action spectra measurements were normalized to 300 µmol m^-2^ s^-1^ after we had confirmed that k_PI_ is directly proportional to light intensity in *E. timida* and *Acetabularia*.

**Supporting information figure S2.**
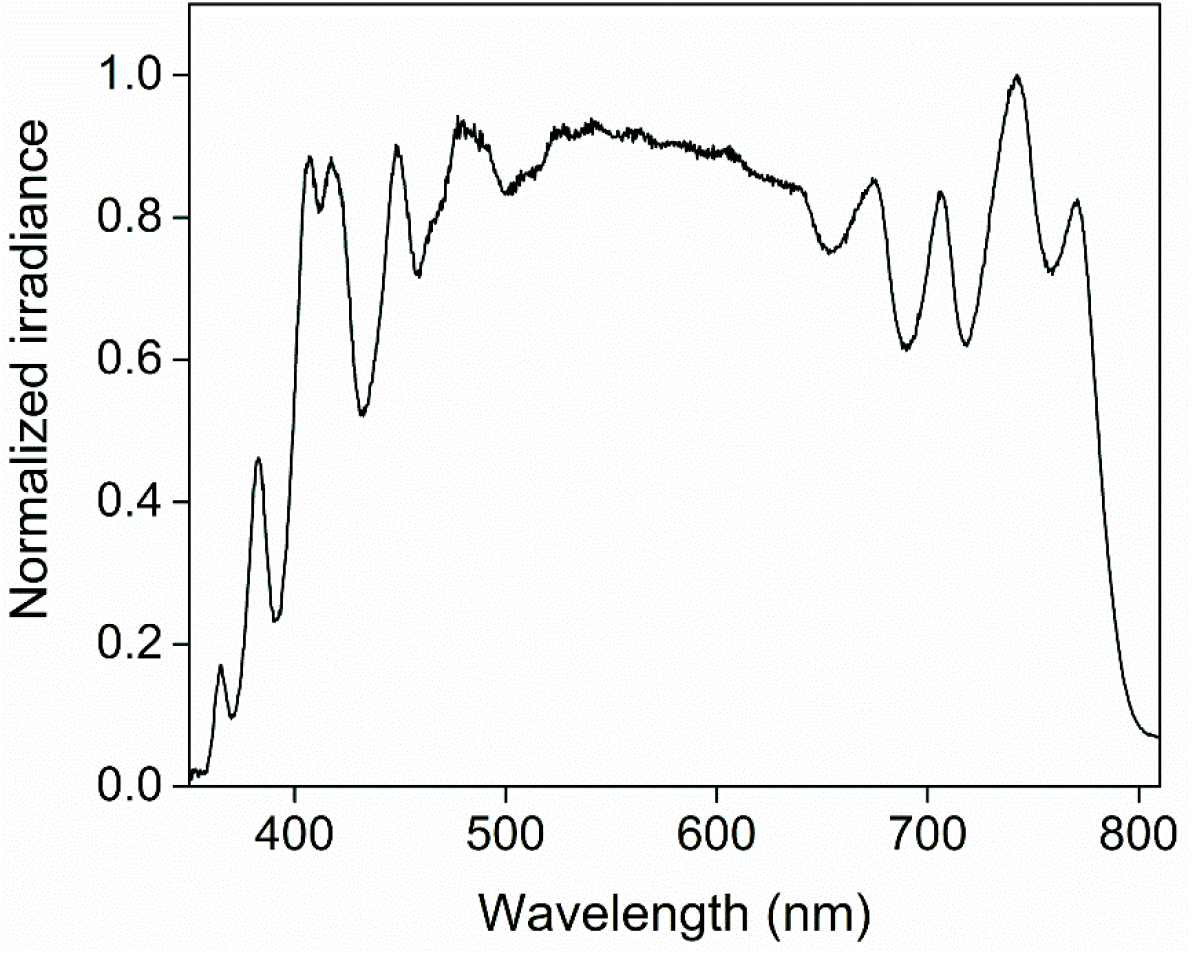
Normalized irradiance spectrum of the Artificial Sunlight Module (SLHolland) that was used as a light source for the white light photoinhibition treatments.

### Room temperature fluorescence emission spectra

For comparison of Chl *a* fluorescence under excitation with different wavelengths, dark acclimated individual slugs or pieces of dark acclimated *Acetabularia* cells were placed on a dry, matte black cardboard and illuminated with low intensity monochromatic light to excite Chl *a*. The algae were cut with a razor blade, and the pieces, amounting to a similar sized clump as an individual slug, were placed in such a manner that the area they covered was similar to that of the slugs. Monochromatic light (450, 470, 490, 510, 530, 550, 590, 600 and 610 nm) was obtained from KL-1500 halogen light sources (Schott AG, Mainz, Germany) filtered through Corion bandpass filters (half width at half maximum 10 nm) via fiber optic light guides. Fluorescence emission excited by 470 nm light was used as a control; the 470 nm excitation light was pointed at the samples at a 45 ° angle, with the head of the light guide fixed to approximately 5 mm away from the sample. Another end of the bifurcated light guide led the emitted fluorescence to the detector of the QE Pro spectrometer (Ocean optics). The light guide used for all other visible light wavelengths was also positioned at 45 ° towards the sample, opposite to the 470 nm light guide.

UV radiation was obtained from a UVA LED (Build My LED; https://www.buildmyled.com/) combined with a 390 nm Corion bandpass filter (full width at half maximum 10 nm). The UV-LED was placed perpendicular to the sample surface and 3 cm away from the sample, so that the end of the spectrometer’s light guide did not obstruct UV radiation. The PFD of the 390 nm excitation was 3 µmol m^-2^ s^-1^, measured with the STS-UV/visible light spectrometer (Ocean optics), whereas the PPFDs of all other wavelengths were 4 µmol m^-2^ s^-1^, measured with a wavelength calibrated PPFD sensor (LI-COR).

For an individual *E. timida* or *Acetabularia* sample, the visible light measurements were carried out by first exciting the sample with 470 nm light and then switching excitation wavelengths (450-610 nm) while maintaining the sample at the exact same position. However, because the UV-excited fluorescence from the slugs was very weak and the correct placement of the samples had to be ensured, the UVA excited fluorescence was always measured first and then the 470 nm excited fluorescence. Fluorescence emission intensities obtained by using different excitation wavelengths were normalized, separately for each individual sample, to fluorescence emission at 685 nm region excited by 470 nm. The normalized fluorescence spectra from all biological replicates were then averaged.

### Confocal microscopy

Individual slugs and *Acetabularia* cells were imaged with an LSM880 confocal with an Axio Observer.Z1 microscope (Zeiss, Oberkochen, Germany) at the Cell Imaging and Cytometry Core, Turku Bioscience Centre, Turku, Finland, with the support of Biocenter Finland. The objective was 20x Zeiss Plan-Apochromat and the acquisition software was ZEN 2.3 SP1. The samples were fixed overnight in the dark at 4 °C with 4 % paraformaldehyde in PBS buffer containing 0.2 % Tween-20. The slugs and the algae were placed in a welled microscope slide. The well was large enough to hold one slug, but the placement of a cover glass over the sample flattened the slug so that the parapodia stayed open. The algae were cut to small pieces to fit in the well. Chl fluorescence was excited with 633 nm light from a HeNe laser and emission was recorded with a GaAsp detector at 640-750 nm range. All images of the slugs were taken from the parapodia, one of the thinnest sections of the slug body, whereas with *Acetabularia* cells pieces of the stalk were imaged. For Z-stacks, the sample was imaged at 3.8 µm intervals by setting the 0 µm layer at a level where Chl fluorescence was still clearly emitted, but nearly out of the focal range. Image analysis was performed with Fiji (Schindelin *et al*., 2012). Maximum intensity fluorescence projections were created using the Z-projection tool, where all slices of the Z-stack contributed to the projection. Average Chl fluorescence of each slice of the Z-stack was obtained by utilizing the “plot Z-axis profile” tool on the entire area of the images without selecting any specific regions of interest. The validity of this method in *Acetabularia*, where the cell area can be accurately defined, was confirmed by comparing the Chl fluorescence of each slice in Z-axis profiles without specific regions of interest to the profiles of Z-stacks where only the *Acetabularia* cell was selected as the region of interest.

### Absorptance and reflectance

Absorptance of intact *Acetabularia* cells and slugs was measured using an integrating sphere (Labsphere, North Sutton, NH, USA). The samples were placed inside a glass test tube in their respective media, and the tube was placed in the integrating sphere. Measurements of the empty sphere were performed with the tubes filled with the media. A 1000 W high pressure Xenon illuminator was used as a light source (Sciencetech Inc., London, Canada) and absorptance was measured with an STS-VIS spectrometer (Ocean optics). The signal-to-noise ratio from the slugs was poor, and this was counteracted by placing 30 live slug individuals in the tube for the measurement. The signal from *Acetabularia* was clear, and approximately 5-10 *Acetabularia* cells were enough to return a sufficient signal for further analysis. All measurements were corrected with the absorptance measurements from absolutely calibrated matte black cardboard (Idle & Proctor, 1983, Pätsikkä *et al*., 1998). However, absolute absorptance values were not calculated because the surface areas of the slugs and *Acetabularia* were unknown. Chls were extracted from the samples by overnight incubation in N,N-dimethylformamide (DMF) in the dark at 4 °C, and the total amounts of Chls were quantified spectrophotometrically using the wavelengths and extinction coefficients for Chls *a* and *b* in DMF (Porra *et al*., 1989).

A nearly identical experimental setup was used for the reflectance measurements as the one used for room temperature fluorescence. Individual slugs or multiple pieces of algae were placed on a matte black cardboard and illuminated with white light from a slide projector guided on to the sample with the bifurcated light guide of the QE Pro spectrometer (Ocean optics). The distance between the probe and the sample was approximately 5 mm. For calibration, a white reflectance standard (Labsphere Inc.) was used to obtain full reflectance using the same setup. In addition to green slugs and *Acetabularia*, spectral reflectance was also measured from starved slug individuals that had lost some of their plastids during starvation.

## Results

### Photoinhibition is slower in *E. timida* than in *Acetabularia*

We measured the decrease in the ratio of variable to maximum fluorescence (F_V_ /F_M_) from both *E. timida* and *Acetabularia* in the presence of lincomycin at seven different light intensities. The decrease in F_V_ /F_M_ in both the slugs and the algae followed first order reaction kinetics (Fig. 1A,B), as usual for photoinhibition of PSII (Tyystjärvi, 2013). In both species, the rate constants of photoinhibition (k_PI_) were directly proportional to light intensity, which indicates that photosynthetic sea slugs are not exempt from this core property of photoinhibition of PSII (Tyystjärvi & Aro, 1996). The k_PI_ values, derived from the measurements in Fig. 1A,B, was approximately twice as high in the algae compared to the slugs in the tested PPFD range (Fig. 1C), indicating that plastids inside *E. timida* are much less prone to photoinhibition of PSII than the plastids inside *Acetabularia* in our experimental conditions.

**Figure 1.**
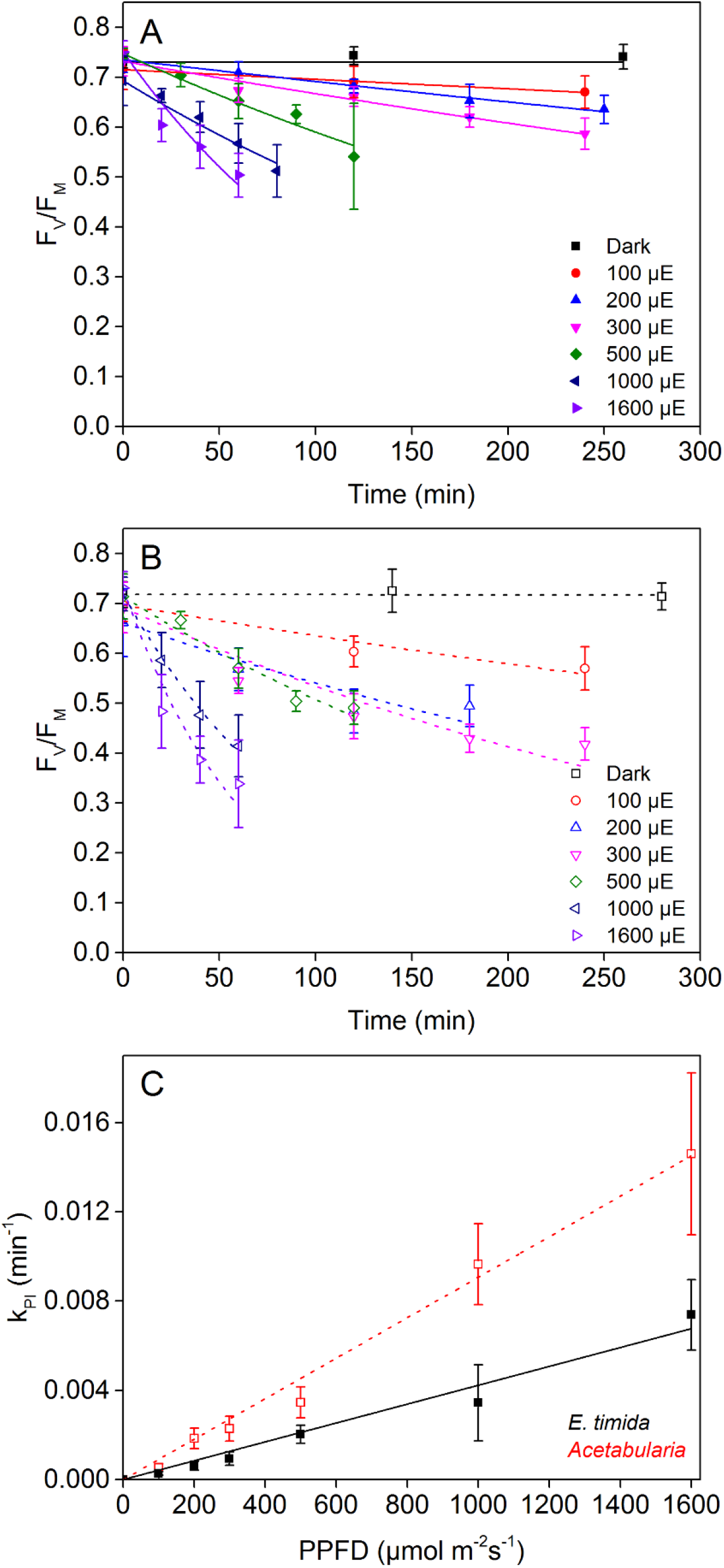
Light response of photoinhibition in lincomycin treated *Acetabularia* and *E. timida*. The decay of the fluorescence parameter F_V_ /F_M_, a proxy of PSII activity, in (A) *E. timida* and (B) *Acetabularia* in response to different PPFDs, as indicated; the designation “µE” stands for µmol of photosyntetically active photons per square meter in a second. The lines show the best fits of the averaged data to first order reaction kinetics (R^2^ of the fits ranged from 0.86 to 0.99 in *Acetabularia* and 0.93 to >0.99 in *E. timida*). (C) Rate constants of photoinhibition (k_PI_) in *Acetabularia* (black) and *E. timida* (red) as a function of PPFD. The lines show linear regression (R^2^ =0.99 and 0.98 for *Acetabularia* and *E. timida*, respectively). k_PI_ values were derived from the measurements shown in (A) and (B). Each data point represents an average of at least four biological replicates and the error bars show SD.

### Slug tissue protects the plastids by screening UV radiation

Next, we measured the action spectrum of photoinhibition from *Acetabularia* and *E. timida*, covering both UV and visible light regions. The results indicate distinct peaks of photoinhibition in both organisms in the visible light region, and UV radiation, compared to visible light, was found to be highly damaging to PSII (Fig. 2). These are common characteristics of photoinhibition shared by all photosynthetic organisms (Jones & Kok, 1966; Havurinne & Tyystjärvi, 2017; Soitamo *et al*., 2017, see Zavafer *et al*., 2015 for review). In both *Acetabularia* and *E. timida* the most pronounced peak in the visible light action spectra is in the red-light region, at 660 nm, but interestingly photoinhibitory efficiency did not significantly drop from 660 to 690 nm. However, the emission spectrum of the 690 nm light shows that our 690 nm source has a contribution of shorter wavelength light that may affect the results (Fig. 2). Other peaks and deeps in the visible light region were more prominent in *E. timida*, where green 560 nm light caused very little photoinhibition, whereas a clear increase in photoinhibition from green to blue 460 and 420 nm light was noticeable. In *Acetabularia*, green 560 and blue 460 nm wavelengths were very similar in their damaging potential.

**Figure 2.**
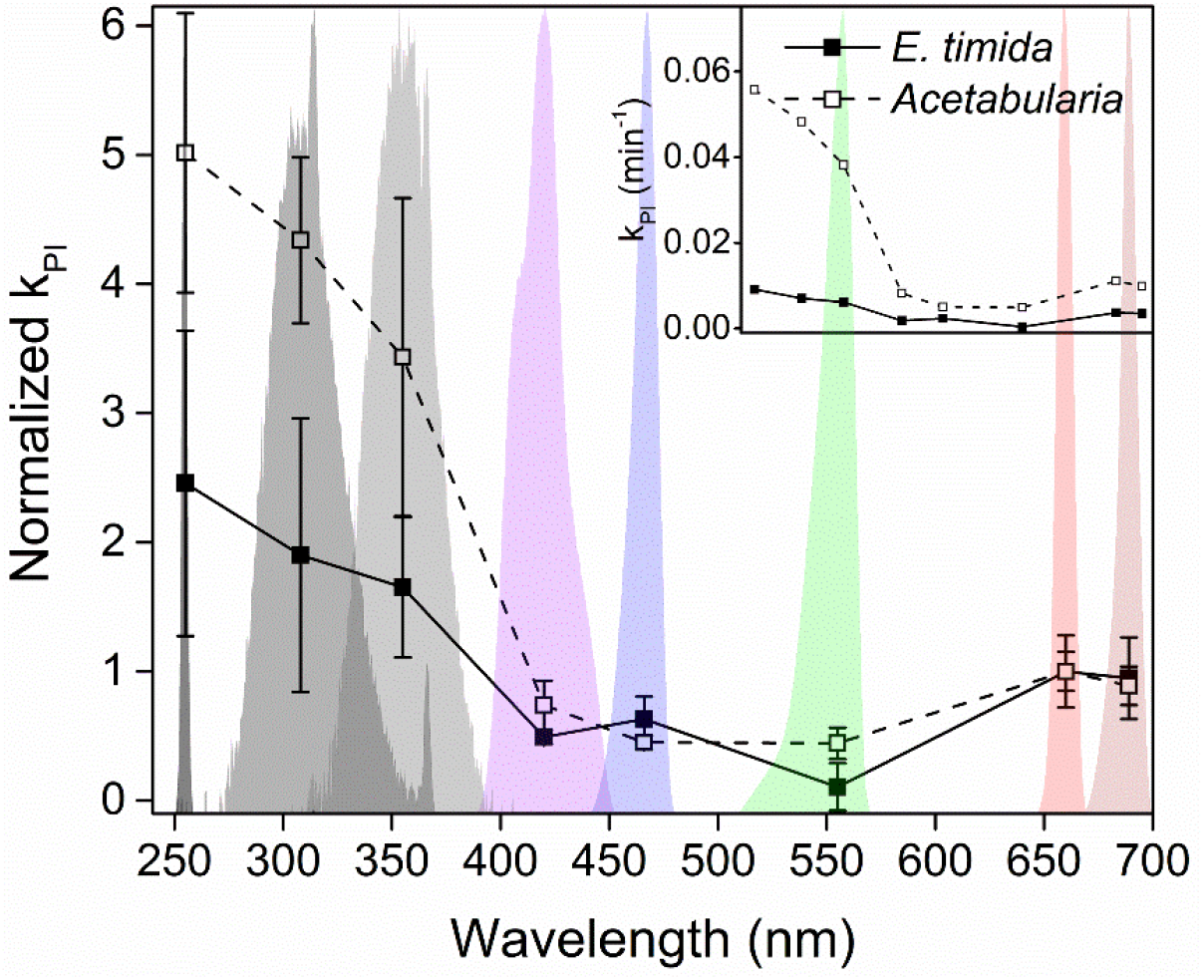
Action spectra of photoinhibition of *Acetabularia* and *E. timida*. The respective treatment light spectra are shown in the background. The rate constants of photoinhibition (k_PI_) have first been normalized to the same (P)PFD (300 µmol m^-2^ s^-1^) and then to the k_PI_ at 660 nm for both species separately to facilitate comparison. The actual treatment light (P)PFDs are detailed in Materials and Methods. The inset shows the action spectra where the k_PI_ values have been normalized only to (P)PFD 300 µmol m^-2^ s^-1^. Each k_PI_ was determined as the best fit to first order reaction kinetics of the decrease in the fluorescence parameter F_V_ /F_M_ during the photoinhibition treatments. Each data point represents an average of at least three biological replicates, and the error bars indicate SD.

Going into the UV region, photoinhibition increased as the wavelength shortened, with UVC causing the most rapid damage in both species. The k_PI_ values of *Acetabularia* and *E. timida* shown in the main Fig. 2 have been normalized to their respective k_PI_ at 660 nm, and they clearly show that UV radiation inflicts very little photoinhibition in the slugs compared to the algae. When the absolute k_PI_ values are compared, the differences are even more dramatic, as the whole action spectrum in *E. timida* seems almost like a flat line in comparison to that of *Acetabularia* (Fig. 2, inset). A slower rate of photoinhibition was already seen in white-light treatments of freshly fed slugs (Fig. 1), but the question remains, why does photoinhibition not increase with decreasing UV wavelength to the same extent in the slugs as in *Acetabularia*?

To elucidate the mechanisms protecting slug plastids against photoinhibition of PSII, we studied the penetration of different wavelengths to the slug tissue. For this, room temperature Chl fluorescence emission at different excitation wavelengths was measured (Fig. 3). Due to the nature of the samples, it was necessary to always perform two measurements (470 nm excitation as a control and another excitation wavelength) from each individual slug or algal mass. With the exception of the UV excitation, the first excitation wavelength was always with 470 nm light to ensure that the samples were in correct position to emit a strong Chl fluorescence signal. For the next measurement from the same sample, the excitation light wavelength was changed to the desired one. This also enabled normalization of the fluorescence emission to the 470-nm-excited fluorescence at 685 nm, facilitating comparison between different samples.

**Figure 3.**
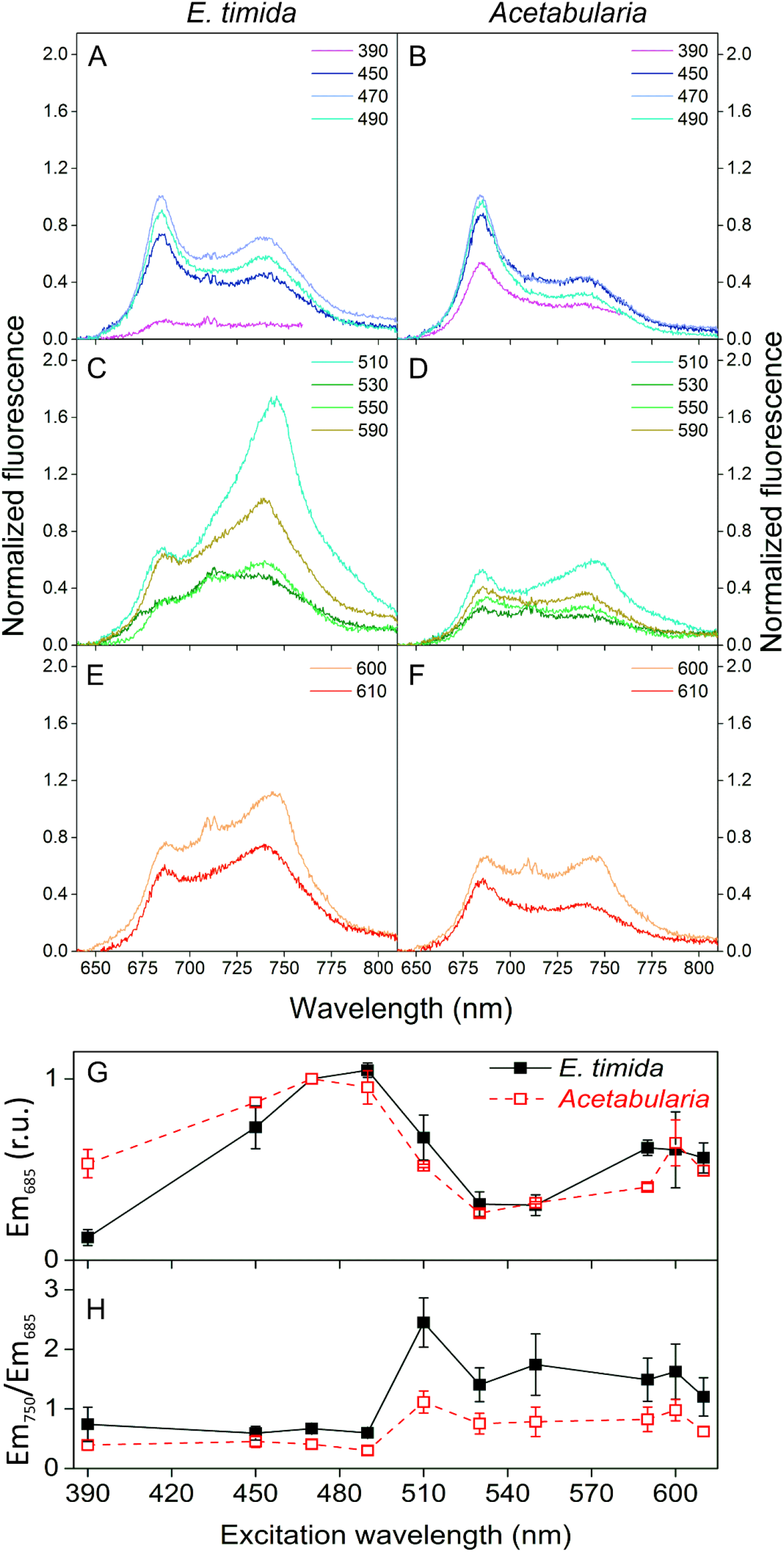
Room temperature Chl fluorescence emission from *E. timida* and *Acetabularia* samples excited with different wavelengths. (A-F) Fluorescence emission spectra from *E. timida* (left panels) and *Acetabularia* (right panels) after excitation with specific wavelengths of light, covering the UV and visible light regions. The excitation wavelengths are indicated in the legends. All fluorescence spectra were normalized to Chl fluorescence emission at 685 nm, excited by 470 nm light. (G) Excitation spectrum of fluorescence emission at 685 nm of *E. timida* (closed symbols) and *Acetabularia* (open symbols), normalized to fluorescence excited by 470 nm light. (H) The ratio of 750 nm to 685 nm fluorescence emission after excitation with different wavelengths. The data in panels G and H were derived from the measurements shown in panels A-F. The PFD of the 390 nm exciting light was 3 µmol m^-2^ s^-1^, whereas for all other wavelengths it was set to 4 µmol m^-2^ s^-1^. Each spectrum and data point represents an average of at least 3 biological replicates, and the error bars indicate SD. The double peak feature at around 710 nm, apparent when the fluorescence signal is low, is a reflected-light artifact.

In the tested UV to blue light excitation wavelengths (390-490 nm), the shapes of the fluorescence emission spectra of *E. timida* and *Acetabularia* showed peaks at the same positions, but the fluorescence emission peak at 750 nm was much more prominent in the slugs (Fig. 3A,B). UV radiation (390 nm) was strikingly less efficient in exciting Chl *a* fluorescence in the slugs than in the algae. In *E. timida* UV-excited fluorescence emission was very weak, whereas in *Acetabularia* the fluorescence yield under UV excitation was only slightly lower than under 470 nm excitation. This indicates that the harmful 390 nm UVA radiation is efficiently blocked from reaching the plastids inside the slugs.

### Fluorescence emission at 685 nm and light absorption per Chl are suppressed in the slugs

The fluorescence excitation spectra of *E. timida* and *Acetabularia* at 685 nm emission wavelength were similar, except for the weaker excitation efficiency of 390 nm in *E. timida* and somewhat weaker efficiency of 590 nm light in *Acetabularia* (Fig. 3G). This indicates that the light-harvesting properties of PSII are similar in both organisms. However, plotting the ratio of 750 nm to 685 nm fluorescence emission against the excitation wavelength revealed strong apparent suppression of 685 nm emission in the slugs (Fig. 3H). Such suppression (apparent enrichment of 750 nm emission) effect can be the result of a higher local concentration of photosynthetic material in *E. timida* in comparison to *Acetabularia*. High local concentration of chlorophyll causes strong self-absorption of fluorescence emission at 685 nm, but not at 750 nm (Lichtenthaler *et al*., 1981; Weis, 1985). Self-absorption also depends on the penetration depth of the excitation light: the deeper the excitation occurs, the higher is the probability of re-absorption of the fluorescence photon on its way out. This feature would also explain why 685 nm fluorescence is strongly suppressed especially in the green excitation wavelength region that is less efficiently absorbed by Chl than blue light (Fig. 3C,H).

The discussion above assumes that the absorption of light in the slugs in the 450 to 610 nm range is dominated by photosynthetic pigments. To test this, we measured both reflectance and absorptance spectra from intact slugs and pieces of *Acetabularia* (Fig. 4). The plastid density inside *E. timida* digestive tubules starts to decrease almost immediately after the slugs are deprived of their food (Laetz *et al*., 2016), which allows for a convenient way of inspecting the effect of plastid abundance on the spectral characteristics of the slugs. The reflectance spectra of freshly fed *E. timida* individuals and *Acetabularia* cells resembled each other, as both reflected far red light (>700 nm) and showed low reflectance in the red and blue regions due to absorption of light by Chl. Reflectance in the green region was higher than in red or blue but lower than in far red, as expected for photosynthetic material (Virtanen *et al*., 2020). Intriguingly, the red edge of reflectance (the increase in reflectance at around 700 nm) in freshly fed slugs appeared to be shifted to longer wavelengths compared to *Acetabularia* (Fig. 4A). The optical properties of *E. timida* changed when the slugs were kept in starvation for 9 and 21 days, or until the slugs were nearly completely bleached, as shown by the reflectance spectra measured from these individuals (Fig. 4A). The difference in the position of the red edge of the reflection spectrum moved toward shorter wavelengths with the proceeding starvation. Interestingly, even the bleached slugs showed a decrease in reflectance with decreasing wavelength from red to blue light, and comparison of the reflection curves of 21 days starved and bleached slugs shows that in blue and green regions the effect of the remaining plastids on reflectance is negligible (Fig. 4A). Thus, the slug tissue absorbs some blue and green light but is transparent to red and far red light.

**Figure 4.**
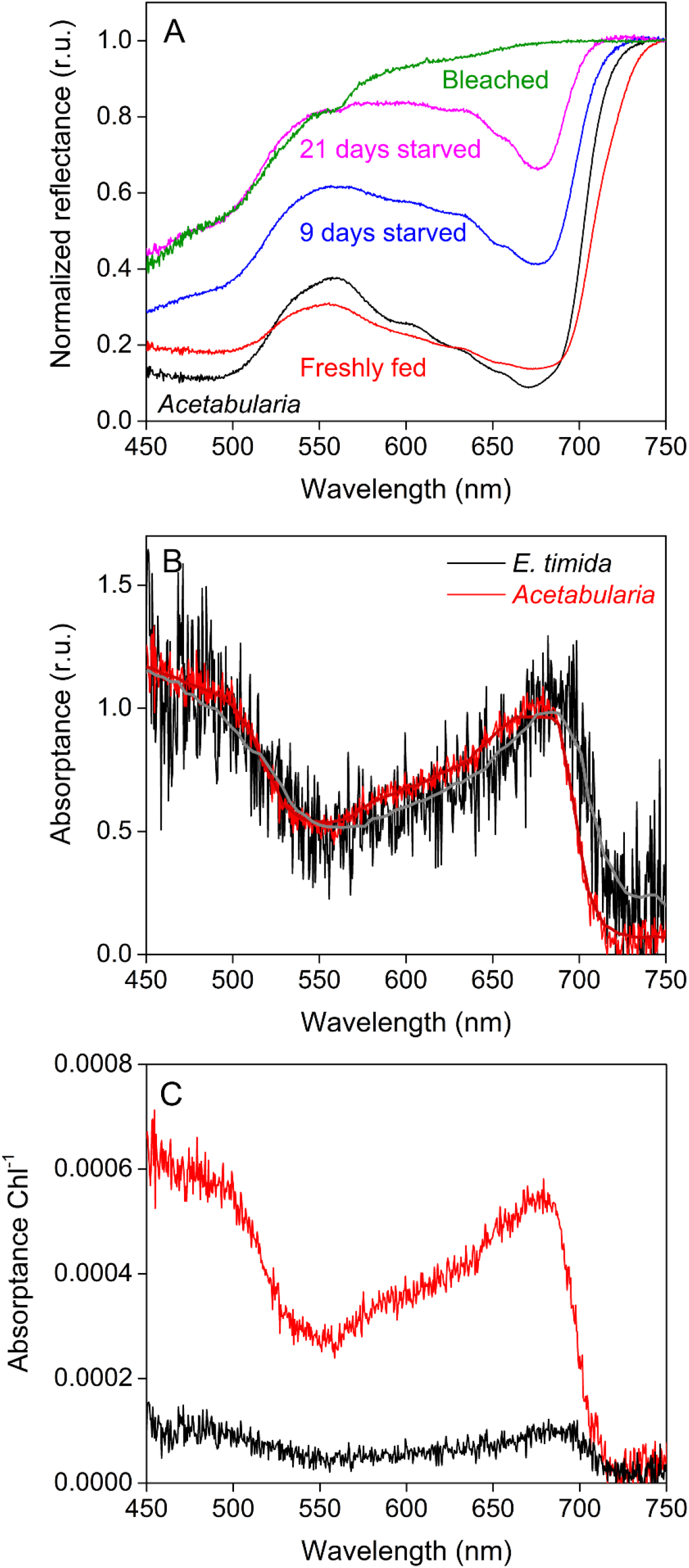
*In vivo* reflectance and absorptance spectra of *Acetabularia* and *E. timida*. (A) Spectral reflectance of *Acetabularia* (black) and *E. timida* individuals that were freshly fed (red), kept in starvation for 9 (blue) and 21 days (magenta), or until the slugs were almost completely bleached and devoid of plastids (green). All reflectance spectra were normalized to their respective reflectance at 750 nm. (B) Absorptance spectra, normalized to the red peaks at around 690 nm and 680 nm for *E. timida* (black) and *Acetabularia* (red), respectively. The bold lines show a running median of the absorptance data. (C) The same spectra as in (B) normalized to the total Chl contents of the samples (87.83 µg Chl for *Acetabularia* and 210.52 µg Chl for *E. timida*). Each curve in (A) represents an average of at least 3 biological replicates. Each curve in (B) and (C) represents an average of three biological replicates for *Acetabularia*, whereas the *E. timida* spectrum represents an average of technical triplicates performed on a sample consisting of 30 slug individuals. Deviations have been omitted for clarity.

The overall shapes of the absorptance spectra of *Acetabularia* and freshly fed *E. timida* were very similar, showing a distinctive red peak (approx. 650-690 nm), low absorptance in the green-yellow region (550-600 nm) and high absorptance in the blue region (450-500 nm) (Fig. 4B). This shape is to be expected for photosynthetic organisms that mainly rely on Chls *a* and *b* for light absorption, such as the green alga *Acetabularia*. However, in accordance with the red shift of the red edge of the reflectance spectra from freshly fed slugs, the red absorptance of the slugs peaked at around 690 nm whereas in *Acetabularia* the red peak was clearly centered around 680 nm (Fig. 4B). Because of the considerably lower signal-to-noise ratio of the slugs compared to the algae (Fig. 4B), artefactual differences cannot be completely ruled out in the absorptance data. Even though the slug tissue itself was found to absorb blue and green light based on the reflectance data (Fig.4A), this seems to be negligible in freshly fed slugs, where the shape of the absorptance spectrum is dominated by photosynthetic pigments, as in *Acetabularia* (Fig. 4B). When the absorptance data were normalized to the total Chl contents of the samples, it became evident that the slugs absorb a lot less light per Chl than the algae (Fig.4C). Although the exact membrane systems surrounding the plastids in *E. timida* are still not resolved, plastids within this slug retain their spherical shape and thylakoid integrity (Wägele *et al*., 2011; Martin *et al*., 2013), suggesting that the lower absorption per Chl in *E. timida* than in *Acetabularia* is likely an indicator of tight, concentrated packing of the plastids in *E. timida* digestive tubules, not a result of changes in the plastid structure.

We also investigated the distribution of plastids inside freshly fed *E. timida* individuals and *Acetabularia* cells using confocal microscopy (Fig. 5). Even though the actual plastid concentration could not be calculated from the micrographs, the images suggest that in the slugs the plastids are arranged in multiple layers within the body, whereas in the algae most of the plastids reside within a narrow layer within the algal cell. Inspecting Chl fluorescence of individual slices of the Z-stack (20 slices spanning 74 µm at intervals of 3.8 µm) revealed that the fluorescence signal in the slugs stayed strong in a wide depth range, whereas the fluorescence signal in *Acetabularia* decreased almost linearly throughout the Z-stack (Fig. 5C).

**Figure 5.**
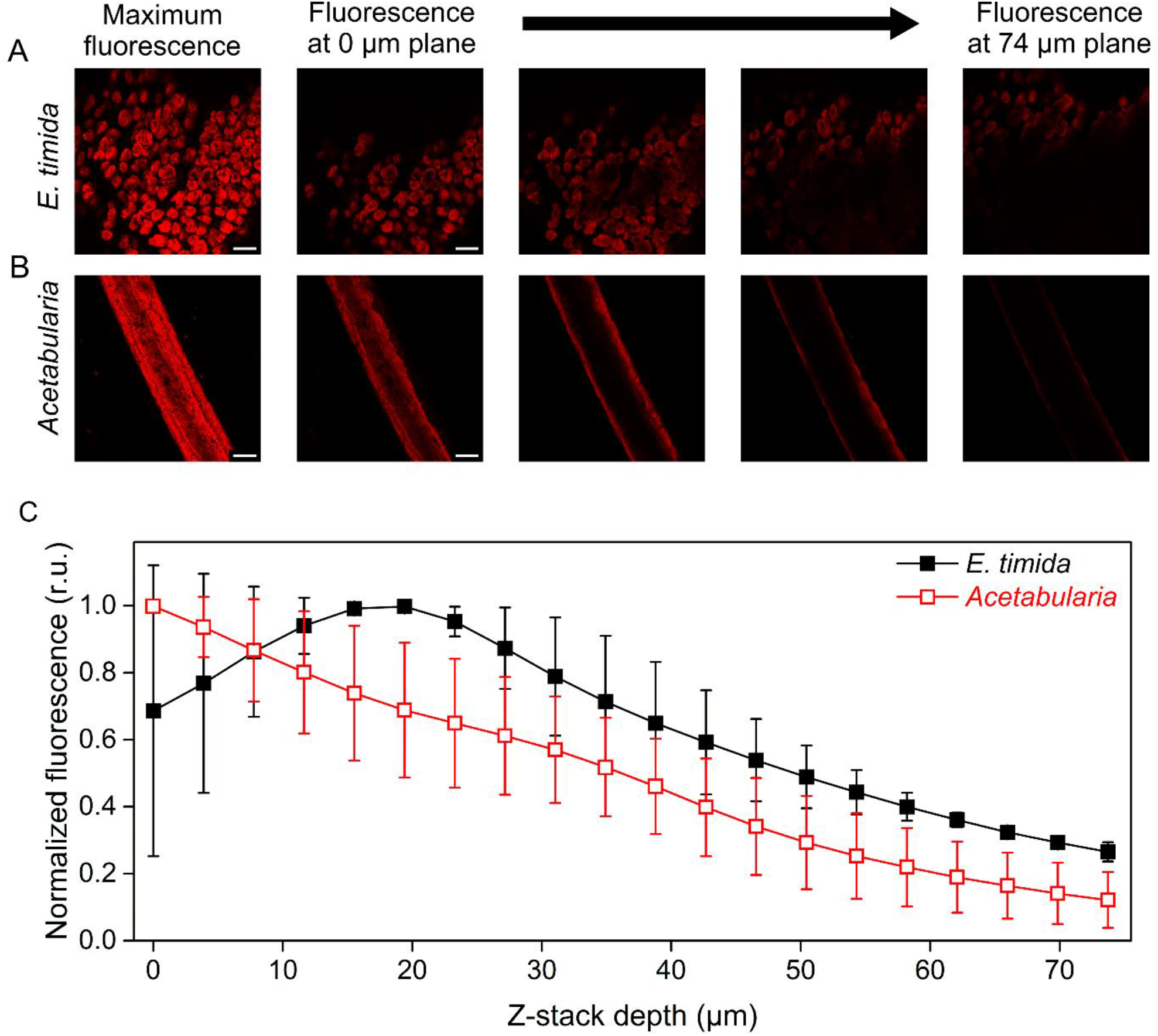
Confocal microscope imaging of Chl fluorescence at different depths inside *E. timida* tissues and *Acetabularia* cells. (A,B) Z-stack images from representative *E. timida* (A) and *Acetabularia* (B) samples. The first images on the left show the projected maximum Chl fluorescence stemming from each individual slice of the Z-stack. The subsequent four-image series (left to right) show Chl fluorescence of individual layers of the Z-stack from the beginning of the stack (0 µm plane) to the end (74 µm plane). The images in between are intermediates at different planes. The scale bars equal 1000 µm. (C) Average Chl fluorescence emission at slices of the Z-stacks at different depths inside the *E. timida* (closed symbols) and *Acetabularia* (open symbols) samples, normalized to their respective maxima. Each data point in (C) represents an average of two biological replicates, and the error bars show SD.

### Starvation makes *E. timida* susceptible to visible-light-induced photoinhibition but has no significant effect on UVA-induced photoinhibition

As the changes in the reflectance spectra of *E. timida* kept in starvation for 9 and 21 days indicated a decrease in the plastid content of the slugs (Fig. 4A), we illuminated these starved slugs in the presence of lincomycin to test if a high plastid concentration protects against photoinhibition. The results show that slugs that had been kept in starvation for 9 days were significantly (P<0.01, n=4, Welch’s t-test) more susceptible to photoinhibition in visible light (PPFD 300 µmol m^-2^ s^-1^) than freshly fed slugs (Fig. 6), and after 21 days the rate constant of photoinhibition (k_PI_ =0.02 min^-1^, SD±0.01, n=7) was approximately 9 times as high as that of freshly fed slugs. The 21-day data may not be equally significant as the 9-day data because the F_V_ /F_M_ of the slugs had already started to decrease during the 21 days of starvation (F_V_ /F_M_ =0.50, SD±0.06, n=7), and their susceptibility to photoinhibition might be affected by a multitude of factors. When 9 days starved slugs were subjected to UVA (365 nm, PFD 33 µmol m^-2^ s^-1^), k_PI_ was not significantly higher than in freshly fed slugs (Fig. 6). The difference between UVA and visible light suggests that a major factor in protecting the plastids against visible light inside the slugs is the high initial plastid concentration in their tissues, whereas the UV protection is caused by the absorption of UV radiation by the slug tissue or mucus.

**Figure 6.**
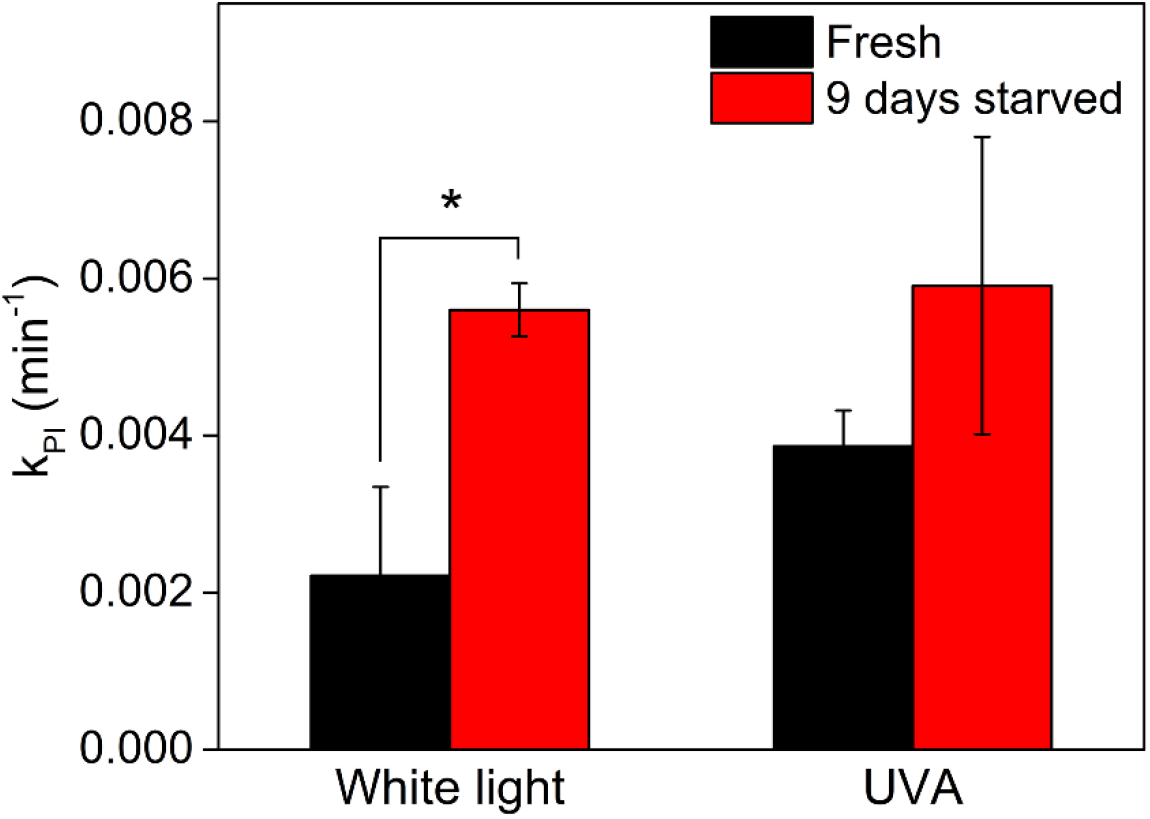
The effect of starvation on susceptibility to photoinhibition in *E. timida*. Rate constant of photoinhibition (k_PI_) induced by white light or UVA, as indicated, of freshly fed *E. timida* slugs (black) and after 9 days in starvation (red). The PPFD of the white light was 300 µmol m^-2^ s^-1^, and the PFD of the UVA radiation treatment was 33 µmol m^-2^ s^-1^. The k_PI_ values from UVA treatments were normalized to PFD 300 µmol m^-2^ s^-1^. A significant difference between the treatments is indicated by an asterisk (*, P<0.01, Welch’s t-test). Each bar represents an average of at least 4 biological replicates and the error bars indicate SD.

## Discussion

### Generalities of photoinhibition hold true for *E. timida*

Three aspects of photoinhibition are nearly ubiquitous among photosynthetic organisms: (I) photoinhibition in the presence of plastid specific translation inhibitors, such as lincomycin, proceeds according to first-order reaction kinetics, (II) k_PI_ is directly proportional to PPFD, and (III) UV radiation causes more damage to PSII than visible light (Tyystjärvi, 2013). All of these “rules” also govern the damage to PSII in the photosynthetic sea slug *E. timida* and its prey, the green alga *Acetabularia* (Figs. 1 and 2). This indicates that the slugs do not alter the fundamental energetic processes of the plastids in a way that would cause deviations to these core properties.

Some photosynthetic slugs, like *E. viridis*, have been shown to either curl up or move to a shadier area when exposed to strong light (Cruz *et al*., 2013; Cartaxana *et al*., 2018), and it was recently shown that also *E. timida* individuals close their parapodia, the wing-like appendices on their sides, in response to increasing light intensity (Cartaxana *et al*., 2019). The authors suggested that this is a photoprotective response. Even though we did not measure the exposed dorsal area of the slugs, we did witness similar behavior during the photoinhibition experiments. However, k_PI_ was directly proportional to PPFD (Fig. 1), although deviation from the linearity would be expected if the shift from open (at PPFD <200 µmol m^-2^ s^-1^ according to Cartaxana *et al*., 2019) to increasingly closed parapodia (PPFD>200 µmol m^-2^ s^-1^) was a major photoprotective measure against photoinhibition of PSII.

### *E. timida* body offers efficient UV sunscreen for its plastids

The majority of studied green macroalgae efficiently repair UV-induced damage rather than screen UV radiation (Pescheck *et al*., 2010; Pescheck *et al*., 2014). Porst *et al*., (1996) showed that *Acetabularia mediterranea* is photoinhibited more by UVB than UVA, and this species recovers from photoinhibition caused by strong natural sunlight almost completely within two hours in the shade. The samples used by Porst *et al*., (1996) were grown naturally in the sea, and therefore likely to have been exposed to at least some UV radiation during their lifetime, which might indicate that also *A. mediterranea* relies on efficient repair instead of UV screening. Likewise, our data show that UVB (and UVC) cause more photoinhibition than UVA in *Acetabularia* (Fig. 2). However, UV-induced photoinhibition is still remarkably slow in *Acetabularia* when compared to plant leaves or diatoms, where UVA causes ten or more times faster photoinhibition than visible wavelengths (Sarvikas *et al*., 2006; Havurinne & Tyystjärvi, 2017). This is intriguing, as the fluorescence emission measurements with UVA excitation reveal that UVA quite efficiently excites Chl in *Acetabularia* (Fig. 4), and thus reaches the plastids.

Sea slugs capable of long-term retention of plastids, including *E. timida*, have genes for a fatty acid synthase-like polyketide synthase (FAS-like PKS), suggesting that they use methylmalonyl-CoA as a substrate to produce polypropionates that can be converted to specific complex polyketides only found in these photosynthetic sea slugs (Torres *et al*., 2020). The presence of complex polyketides in *E. timida* has also been confirmed, and the compounds identified in it include *ent*-9,10-deoxytridachione, tridachione, photodeoxytridachione and 15-norphotodeoxytridachione (Gavagnin *et al*., 1994; Torres *et al*., 2020). These compounds have been suggested to function as UV sunscreens in the mucus excreted by the slugs (Ireland & Scheuer, 1979), and they are related to the plastids, as the mucus contains a large fraction of the radiolabeled carbon originating from carbon fixation by the slug plastids (Trench *et al*., 1972). Accordingly, our results show that the slug tissue efficiently blocks UV radiation from reaching the plastids (Fig. 3) thereby protecting the plastids from UV-induced photoinhibition (Fig. 2). The protection appears to function even in starved slug individuals, indicating that this protection mechanism is different from the mechanism that protects in visible light (Fig. 6). Screening of UV radiation is expected to have a significant effect on plastid longevity in *E. timida*, as sunlight has a considerable UVA contribution and UVA is efficient in causing photoinhibition (Fig. 2). However, our data do not allow us to identify the screening molecules.

If *Acetabularia* itself has evolved to deal with UV radiation by efficient repair, then plastids in *E. timida* would benefit from both efficient repair machinery of the alga (at least to the extent that is possible without the algal nucleus) and UV screening of the slug. While the genetic autonomy and efficient repair machinery of *Vaucheria litorea* plastids are likely a major factor in maintaining the plastids functional in the sea slug *E. chlorotica* (Green *et al*., 2000; Havurinne *et al*., 2021), there are contrasting reports on the capability of plastids inside *E. timida* to recover from photoinhibition. Whereas Christa *et al*., (2018) found no difference in the repair of photodamage in *E. timida* in the presence or absence of lincomycin after a 30 min recovery period following a 1 h photoinhibition treatment, our previous results suggest otherwise; when the slugs were allowed to recover overnight after photoinhibition, lincomycin strongly inhibited the recovery (Havurinne *et al*., 2021). We do agree with the statement made in Christa *et al*., (2018) that the plastids inside *E. timida* do not recover as efficiently as they do inside *Acetabularia* (Supporting Information Fig. S1), but the data in Havurinne *et al*., (2021) show that the inherent repair machinery of the plastids does likely play a role in maintaining the plastids functional also in *E. timida*.

### Tight packing of plastids within *E. timida* protects from photoinhibition

*Acetabularia* cells appear transparently green whereas the areas with plastids in the slugs are bright green. The spectral characteristics of freshly fed *E. timida* and *Acetabularia* corroborate these ocular observations and show that the plastids of *E. timida* are tightly packed, in comparison to the plastids within their original host. Firstly, *E. timida* exhibited a strong suppression of 685 nm fluorescence due to self-absorption compared to *Acetabularia* in the tested excitation wavelength range (Fig. 3). Reflectance and absorptance spectra provide a second piece of evidence for tight packing of plastids within the slugs, as the red edge of reflectance and the red absorption peak of Chl *a* in freshly fed *E. timida* are shifted in a manner that suggests that the slugs absorb longer wavelengths of red and far-red light than *Acetabularia* (Fig. 4A,B). The same phenomenon can be seen in senescing birch leaves, as green leaves absorb light at longer wavelengths than senescing ones (Mattila *et al*., 2021). Furthermore, the red edge of the reflectance spectrum of *E. timida* moves toward shorter wavelengths when the slugs lose plastids during starvation (Fig. 4A). These data show that the position of the red edge of reflectance, and consequently also that of the red peak of absorptance, depend on the amount of green plastids per slug. Tight packing of slug plastids also explains why the slugs absorb much less light than the algae when the absorptance is normalized to the Chl content of the samples (Fig. 4C). It should be noted, however, that 30 slugs had to be packed in a test-tube to get a single absorptance reading, and therefore the packing of slugs may have further lowered the absorptance. Results of confocal microscopy further confirm that plastids inside *E. timida* are spread to a wider depth range than plastids in *Acetabularia* (Fig. 5).

Tight packing of plastids inside the slug tissue can explain why the slug plastids appear to be less prone to photoinhibition than the same plastids in *Acetabularia* (Fig. 1; Christa *et al*., 2018). The mechanism is simple: the outermost plastids of a tight stack prevent light from reaching the lower ones. Protection against photoinhibition by a high Chl concentration was shown in plant leaves by Pätsikkä *et al*., (1998), and methods to model the intrinsic rate constant of photoinhibition in optically thick samples have recently been elaborated (Serôdio *et al*., 2014; Serôdio & Campbell, 2021). The finding that the susceptibility of algal plastids to visible-light photoinhibition increases when plastids are lost during starvation (Fig. 6) confirms that the packing of plastids protects their PSII against photoinhibition.

The finding that tight packing of plastids is a major mechanism of protection against photoinhibition of PSII does not exclude possible protection by other mechanisms. Plastids inside *E. timida* maintain physiological photoprotection mechanisms, such as the xanthophyll cycle (Christa *et al*., 2018; Cartaxana *et al*., 2019). However, the effect of NPQ on photoinhibition of PSII is usually small (Tyystjärvi, 2013), and other photoprotective mechanisms found in slug plastids (Havurinne & Tyystjärvi, 2020) would only marginally protect PSII. Our data suggest that, if given the chance, *E. timida* slugs fill their thick bodies up with plastids, which protects the plastids from photoinhibition of PSII. Slow photoinhibition improves the longevity of the plastids inside photosynthetic sea slugs.

Photosynthetic sea slugs can move away from excessive irradiation if the need arises. Nevertheless, periods of strong light are inevitable in the shallow waters that slugs like *E. timida* inhabit. Our results demonstrate that the slugs protect their plastids by screening highly damaging UV radiation and by packing their plastids tightly all over their bodies, allowing the outer layers to take the brunt of the damage.

## Acknowledgements

This work was supported by Academy of Finland (grant 333421, to ET). VH would like to thank Finnish Cultural Foundation, Finnish Academy of Science and Letters, Turku University Foundation, University of Turku Graduate School and Kone Foundation for financial support. HM would like to thank the Emil Aaltonen Foundation. Iiris Kuusisto is thanked for building the custom LED system that was used in the photoinhibition experiments.

## Author contributions

VH and ET conceptualized the study. RA and VK performed most of the photoinhibition experiments and were involved with culturing *E. timida* and *Acetabularia*. HM aided in experimental design, measured the absorptance spectra and carried out part of the photoinhibition experiments. VH did the reflectance and fluorescence measurements and wrote the first draft of the manuscript. All authors contributed to finalizing the manuscript. ET supervised the study.

## Supporting information

Supporting information figure S1 – The effect of different lincomycin concentrations to the recovery of PSII activity in *Acetabularia* after photoinhibition.

Supporting information figure S2 – The normalized irradiance spectrum of the white light lamp used in polychromatic white light photoinhibition experiments.

